# Functional Diversity in GII.4 Norovirus Entry: HBGA Binding and Capsid Clustering Dynamics

**DOI:** 10.1101/2025.05.20.655000

**Authors:** B. Vijayalakshmi Ayyar, Carmen V. Apostol, Janam Dave, Soni Kaundal, Frederick H. Neill, Khalil Ettayebi, Sarah Maher, Ramakrishnan Anish, Göran Larson, Robert L. Atmar, Sue E. Crawford, B. V. Venkataram Prasad, Mary K. Estes

## Abstract

Human noroviruses (HuNoVs), especially GII.4 strains, are the leading cause of acute viral gastroenteritis worldwide, yet no approved vaccines or antivirals exist. The pandemic GII.4 Sydney 2012 strain enters cells via membrane wounding and clathrin-independent carrier (CLIC)-mediated endocytosis, but it is unclear whether this entry mechanism is conserved across GII.4 variants.

We compared early binding and entry of multiple GII.4 variants using wildtype and mutant GII.4 virus-like particles (VLPs) and modified human intestinal enteroid (HIE) cultures. Only a subset of GII.4 variants, including GII.4 Sydney, form distinct, HBGA-dependent capsid clusters on the cell surface. Clustering strains display significantly enhanced membrane wounding and endocytosis compared to non-clustering strains and outcompete non-clustering strains in replication assays as shown by complete inhibition of GII.4 Sydney replication.

Using mutant VLPs and a HBGA non-binding mutant (R345A), we identified two residues, V333 and R339, in the VP1 protruding domain as critical mediators of clustering and entry. Mutations of these residues disrupt clustering and endocytosis without affecting HBGA binding, suggesting a role in post-attachment processes. While clustering and endocytosis are contingent upon VLP binding to HBGAs, inhibitor studies show they are independent of host protein glycosylation and are driven by lipid raft remodeling regulated by cholesterol and ceramides.

Quantitative analyses across multiple GII.4 variants reveal an apparent dichotomy between clustering and non-clustering phenotypes, with clustering variants exhibiting higher entry competence. This distinction offers insight into strain-specific cell entry mechanisms and may aid in identifying the elusive proteinaceous HuNoV cellular receptor(s) supporting targeted therapeutic development.

## Introduction

Human noroviruses (HuNoVs) are the leading cause of acute viral gastroenteritis worldwide, yet effective vaccines and therapies remain elusive. These viruses are highly infectious and genetically diverse, with 37 identified genotypes based on capsid protein sequences (Chhabra et al., 2019). Among these, over the past two decades the GII.4 genotype has been the most prevalent, being responsible for at least six pandemics (White, 2014). Remarkably, despite continual antigenic drift and the emergence of new GII.4 variants, the Sydney strain persists as a dominant circulating lineage—a testament to its superior infection capabilities, environmental stability, and rapid spread (Carlson et al., 2024; Prasad et al., 2025).

Viral entry, the first stage of the viral life cycle, is a critical determinant of cell tropism, host range, and pathogenesis. Successful HuNoV replication requires expression of the correct carbohydrate histo-blood group antigens (HBGAs) needed for initial virus binding to the cell. The HBGA binding capsid protein VP1 protruding (P-) domain is under strong immune selection, driving capsid changes that balance receptor binding with immune escape (Lindesmith et al., 2022; Lindesmith et al., 2008). The determination of how these capsid adaptations optimize entry efficiency may contribute to understanding their role in the persistence of pandemic GII.4 variants.

Our recent research uncovered that the pandemic GII.4 Sydney 2012 uses a sophisticated entry mechanism to infect human intestinal enteroids (HIEs). This process involves a complex interplay between membrane wound repair and clathrin-independent carriers (CLIC)-mediated endocytosis. GII.4 Sydney binds fucosylated HBGAs on secretor-positive enterocytes, triggering focal membrane damage that recruits galectin-3 (Gal-3) and lysosomal-associated membrane protein 1 (LAMP-1) to the plasma membrane, primes CLIC-mediated uptake, and drives infection (Ayyar et al., 2023). While core HBGA contacts in the viral VP1 P-domain are conserved, adjacent residues fine-tune affinity and tropism (De Rougemont et al., 2011; Esseili et al., 2019; Liang et al., 2021). In addition to HBGA binding, bile acids are obligatory for replication of some HuNoV strains (GII.3, GI.1, GII.6, and GII.17) compared to GII.4, which do not require bile acid for entry and replication. Interestingly, bile acid-dependent HuNoV strains do not induce endocytosis. These observations raise questions about the universality of CLIC-mediated endocytosis for GII.4 variants, the role of strain-dependent variations in HBGA affinity, and the differential effects of bile acids in entry and infectivity of HuNoVs (GII.4) that only infect secretor-positive HIEs or strains (GII.3) that can infect both secretor-positive and secretor-negative HIEs (Ayyar et al., 2023; Ettayebi et al., 2016; Murakami et al., 2020).

In this study, using virus-like particles (VLPs), we found that GII.4 Sydney entry relies on VP1-driven, HBGA-mediated nanoscale clustering of capsids at the HIE apical membrane. With other viruses, clusters form high-avidity platforms that enhance membrane curvature and membrane fusion efficiency, increase the multiplicity of infection, and shield virions from immune detection (Andreu-Moreno & Sanjuán, 2018; Helenius, 2018). Thus, clustering is not just a structural phenomenon but an important process for viral uptake and replication (Cureton et al., 2012; Ewers et al., 2010; Sieben et al., 2020; Wong et al., 2021). For non-enveloped viruses, this mechanism mirrors simian virus 40 (SV40), which binds to GM1 ganglioside receptors to drive CLIC-like uptake into glycolipid-deficient murine myeloma GM95 cells (Ewers et al., 2010). Similarly, human papillomavirus (HPV16) clusters heparan sulfate for endocytosis into HeLa cells (Spoden et al., 2008). Both viruses utilize clathrin-independent endocytosis mechanisms for cell entry, highlighting a conserved viral paradigm where glycan/lipid-driven capsid clustering promotes viral internalization.

Herein, comparing binding, clustering and uptake of bile acid–dependent versus independent HuNoV genotypes, multiple GII.4 variants, as well as wild-type (WT) and mutant VLPs, we report how subtle variations in VP1 dictate membrane engagement, clustering efficiency, and entry dynamics. These insights increase our understanding of norovirus cell entry mechanisms and lay the groundwork for identifying the elusive HuNoV receptor and guiding the development of targeted therapeutic interventions.

## Results

### GII.4 Sydney triggers receptor clustering and activates CLIC-mediated endocytosis in secretor-positive HIEs

CLIC-mediated endocytosis is initiated by glycan-dependent oligomerization of cell surface receptors, leading to lateral clustering and membrane deformation that generates tubular carriers (Lakshminarayan et al., 2014). Our previous work showed that pandemic GII.4 Sydney activates this pathway in J2 HIEs by engaging initial glycosylated apical binding factors, promoting actin remodeling and membrane invagination (Ayyar et al., 2023). To investigate early viral uptake, we first performed confocal and time-lapse microscopy on GII.4 Sydney-VLP inoculated HIEs and observed the formation of discrete VP1 capsid protein clusters (>3-5 pixel^2^) on the apical surface (Fig. 1a, b). Co-staining with the glycosylated apical membrane protein ACE2 confirmed that these clusters localize to the apical domain and increase over time, consistent with progressive receptor clustering during CLIC-dependent internalization (Fig. 1c).

**Figure 1.**
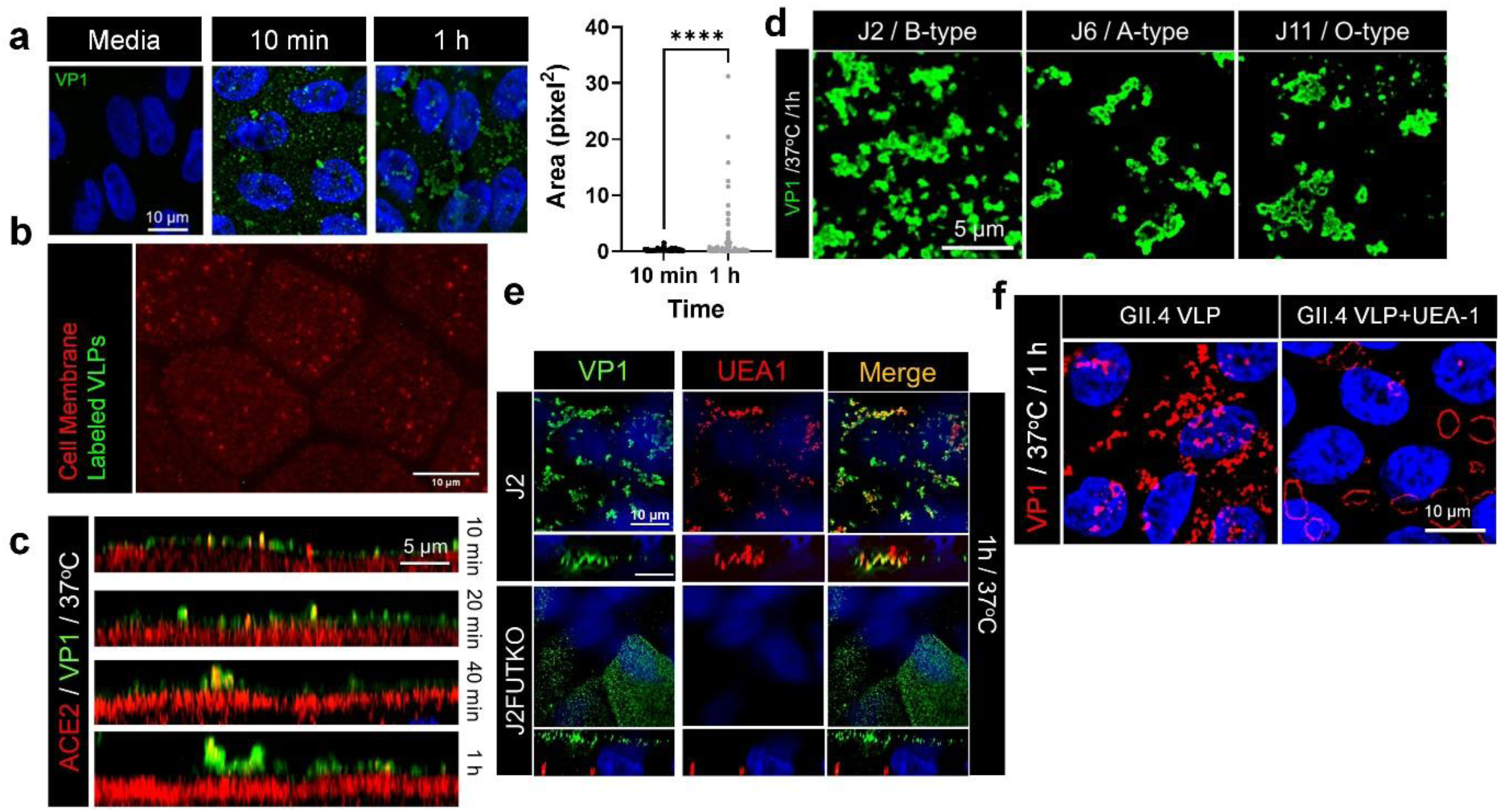
GII.4 Sydney capsid clustering occurs apically in secretor positive HIEs regulated by *FUT*2 expression. **(a)** Confocal microscopy showing binding of GII.4 Sydney VLPs to secretor positive J2 HIEs at 10 minutes (dispersed) and 1 h (clustered) at 37oC. Inset: quantification of area showing VP1 staining at 10 min and 1 h of VLP incubation at 37oC representing clustering (1 pixel = 0.04 μm) Significance (P values) calculated using unpaired t test with Welch’s correction. **(b)** Time lapse microscopy showing GII.4 Sydney VLP cluster formation (green) over time (1 h at 37oC) on membrane surface (red) using Alexa Fluor labelled GII.4 Sydney VLPs. **(c)** ACE2 staining of HIE membranes showing GII.4 Sydney capsid clustering (VP1) occurs on HIE apical surface at 37oC. **(d)** Confocal microscopy showing GII.4 Sydney VLP cluster formation in secretor positive HIEs occurs in HIEs expressing all ABO glycans. **(e)** Confocal microscopy showing colocalization of GII.4 Sydney VP1 (green) and UEA-1 in (red) secretor positive J2 HIEs compared to isogenic secretor negative J2FUT2KO HIEs. **(f)** Pre-incubation of J2 HIEs with UEA-1 inhibits clustering of GII-4 Sydney VLP (red) on the plasma membrane surface measured post 1 h incubation at 37oC.

GII.4 Sydney-induced clustering (>3 pixel^2^) was detected in secretor-positive HIEs from multiple donors, from all ABO blood group genotypes (Fig. 1d; Table 1), but was abolished in *Fucosytransferase 2* (*FUT2*)-deficient lines (Supplementary Fig. 1a), indicating a requirement for α1,2-fucosylated glycans. Clustering was restricted to small intestinal HIEs, with no detectable clustering in non-permissive colonic lines (Supplementary Fig. 1b, c; Ettayebi et al., 2024).

**Table 1:** HIE characteristics.

| Table 1: HIE characteristics |  |  |  |  |  |
| --- | --- | --- | --- | --- | --- |
| HIE line | Source age | Phenotype |  |  | GII.4 replication |
|  |  | Secretor status | ABH Status | Lewis status |  |
| J2 | 52 y | Positive | B | Le<sup>b</sup> | + |
| J2FUT2KO | 52 y | Negative | B | Le<sup>a</sup> | - |
| J4 | 62 y | Negative | O | Le<sup>a</sup> | - |
| J6 | 43 y | Positive | A | Le<sup>b</sup> | + |
| J8 | 52 y | Negative | O | Le<sup>a</sup> | - |
| J11 | NA | Positive | O | Le<sup>b</sup> | + |
| 2005 (D,J,I) | 23 y | Positive | O | Le<sup>b</sup> | + |
| C2005 | 23 y | Positive | O | Le<sup>b</sup> | - |
| All the tested HIEs are Lewis positive |  |  |  |  |  |

To further probe the role of fucosylated glycans in clustering, secretor-positive HIEs were stained with UEA1, a lectin that binds terminal α1,2-fucose. This revealed fucosylated glycan clusters co-localizing with VLPs (Fig. 1e). UEA1 preincubation of HIEs reduced VLP-induced clustering and changed the membrane distribution pattern of the VLPs (Fig. 1f), suggesting competition for and redistribution of fucosylated receptor sites (Parveen et al., 2019). Functionally, UEA1 pretreatment before infection significantly reduced GII.4 replication despite increased initial viral binding (Supplementary Fig. 1d), confirming that receptor clustering, rather than binding alone, is essential for viral entry and productive infection.

### Secretor status dictates GII.4 entry via glycan-mediated signaling and CLIC pathway activation

It is well established that GII.4 binding to HBGAs is necessary for infection (Haga et al., 2020). To further investigate the role of HBGAs in viral entry and downstream signaling, we compared GII.4 Sydney VLP interactions in the isogenic J2 HIE line and its genetically modified J2*FUT*2-knockout (J2FUT2-KO) line, in which the *FUT2* gene is disrupted, resulting in the absence of surface expression of α1,2-fucosylated HBGAs (Haga et al., 2020). Confocal microscopy (Fig. 2a), Western blotting of membrane-associated VLPs (Fig. 2b), and binding assays (Fig. 2c) confirmed significantly reduced GII.4 attachment in J2FUT2KO cells. However, residual VLP binding still persisted in this secretor negative line (Supplementary Fig. 1a), suggesting that *FUT*2-independent alternative surface molecules such as GalCer (Bally et al., 2012) or Lewis glycans may contribute to GII.4 binding (Tarris et al., 2022).

**Figure 2.**
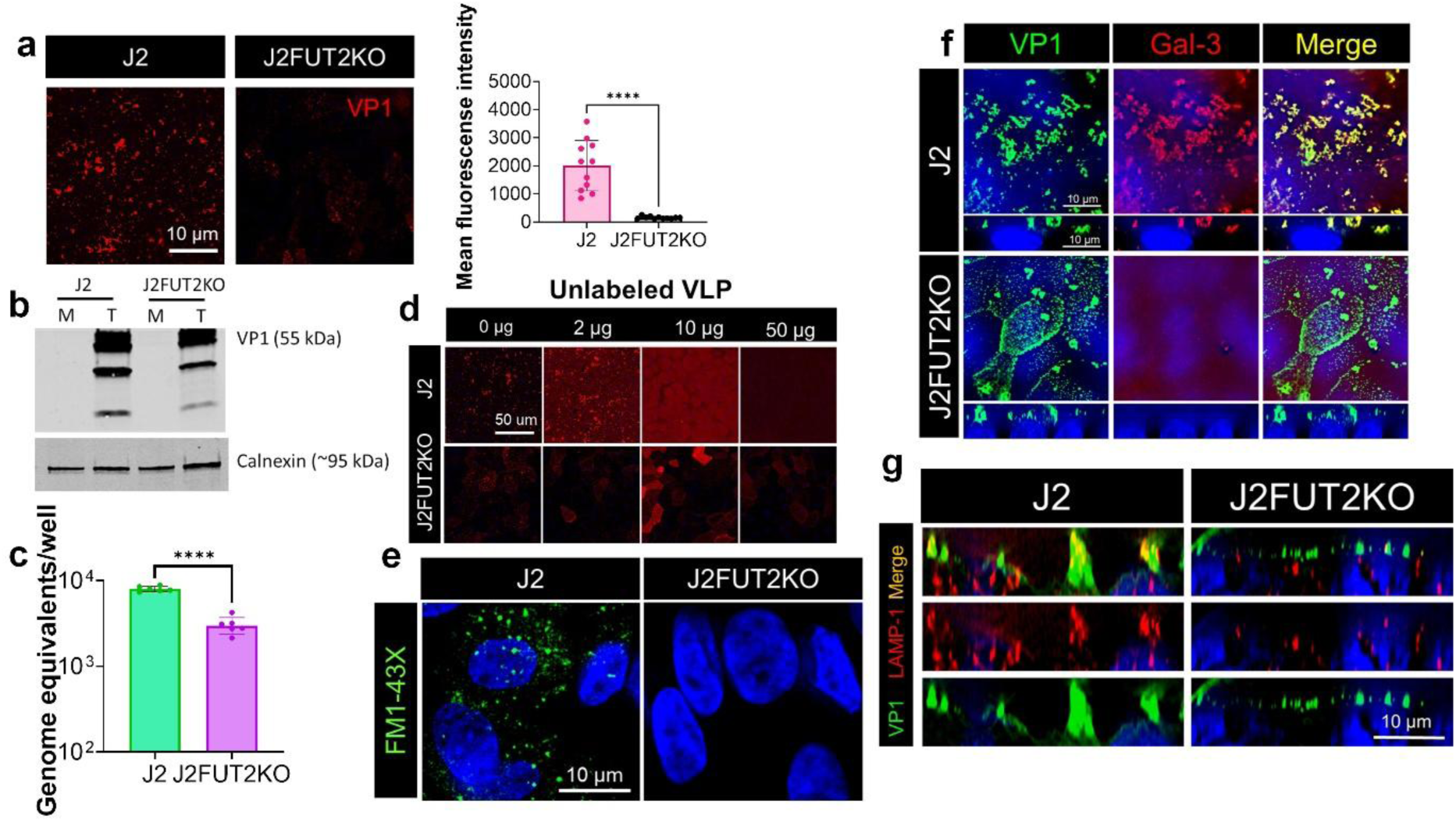
Maximal GII.4-induced binding and subsequent virus-induced endocytosis and lysosomal exocytosis only occur in secretor positive HIEs. **(a)** Confocal microscopy showing binding of Alexa Fluor labelled-GII.4 Sydney VLPs (red) following 1 h incubation at 4°C to secretor positive J2 and isogenic secretor negative J2FUT2KO HIEs along with its quantification (right). Significance (P values) calculated using unpaired t test with Welch’s correction. **(b)** Western blot analysis showing GII.4 Sydney VLP binding (VLP) to proteins in total membrane preparations isolated following 1 h incubation of VLPs at 37°C from J2 and J2FUT2KO HIEs. M and T show proteins in membrane preparations from media and VLP-treated cells. Calnexin was used as a loading control. The full-length capsid protein and cleavage products were detected. **(c)** RT-PCR quantification of GII.4 Sydney virus (from stool) bound to J2 and J2FUT2KO HIEs 1h post-inoculation at 37°C (n=2). **(d)** Inhibition assay showing binding at 4°C of Alexa Fluor labelled GII.4 Sydney VLPs (red) in the presence of increasing concentrations of unlabeled VLPs in J2 and J2FUT2KO HIEs. **(e)** FM-1-43FX uptake detecting endocytosis induced 10 minutes after treatment with unlabeled GII.4 Sydney VLPs and incubation at 37°C in J2 and J2FUT2KO HIEs. **(f)** Confocal microscopy comparing GII.4-induced Gal-3 recruitment to the apical membrane surface and its colocalization with GII.4 Sydney VP1 in both J2 and J2FUT2KO HIE lines. **(g)** GII.4-induced lysosomal exocytosis detected by LAMP-1 staining at the plasma membrane observed at 1h at 37°C after GII.4 VLP inoculation only in J2HIEs and not in J2FUT2KO HIEs.

To further assess binding specificity, we performed a competition assay using unlabeled GII.4 Sydney VLPs to compete with fluorescently labeled VLPs. Unexpectedly, at low competitor concentrations, VLP binding increased in both J2 and J2FUT2KO cells, while high competitor concentrations inhibited binding completely (Fig. 2d) suggesting the possibility that there are optimal concentrations for different kinds of clustering and to different receptors (Parveen et al., 2019). Maximal binding was observed for incubations with unlabeled VLP at 2 µg in J2 and 10 µg in J2FUT2-KO cells, indicating that GII.4 binding is specific and saturable in both cell types. Despite some detectable binding in J2FUT2KO cells, GII.4 endocytosis was observed only in J2 cells and not in J2FUT2KO HIEs (Fig. 2e). To further probe the role of VLP binding to HBGAs in endocytosis, we examined the apical redistribution of Gal-3, and LAMP-1.

Gal-3 is a CLIC pathway regulator linked to glycan-dependent signaling, and LAMP-1 is a marker of apical lysosomal exocytosis. Upon VLP exposure, both Gal-3 and LAMP-1 were mobilized to the apical surface in J2 cells but not in J2FUT2KO cells (Fig. 2f, g). These data, together with previous findings (Ayyar et al., 2023), suggest that initial virus–host interactions can trigger host signaling pathways that facilitate downstream events such as receptor trafficking and endocytosis.

### GII.4 clustering and viral entry are dependent on lipid rafts and cholesterol-mediated membrane dynamics

As noted earlier, CLIC-mediated endocytosis is tightly regulated by lipid raft microdomains enriched in cholesterol and glycolipids, which organize membrane curvature and cargo clustering (Ivashenka et al., 2021; Lakshminarayan et al., 2014). To investigate whether GII.4 capsid clustering is raft-dependent, we incubated HIEs with GII.4 Sydney VLPs and isolated detergent-resistant membranes. Western blotting revealed VP1 enrichment in lipid raft fraction, containing flotillin, a raft-resident marker (Fig. 3a), suggesting that the viral capsid partitions into rafts during the initial stages of entry. To assess whether viral engagement alters raft dynamics, we used confocal microscopy to monitor phosphatidylinositol 4,5-bisphosphate (PIP2), a key lipid that promotes protein sorting and nanocluster formation within raft-associated regions (Katan & Cockcroft, 2020). A time-dependent increase in PIP2 intensity was observed at the apical surface of J2 HIEs, but not in J2FUT2KO cells (Fig. 3b), indicating that HBGA-dependent binding reorganizes membrane lipids to initiate capsid clustering and recruitment into the CLIC pathway.

**Figure 3.**
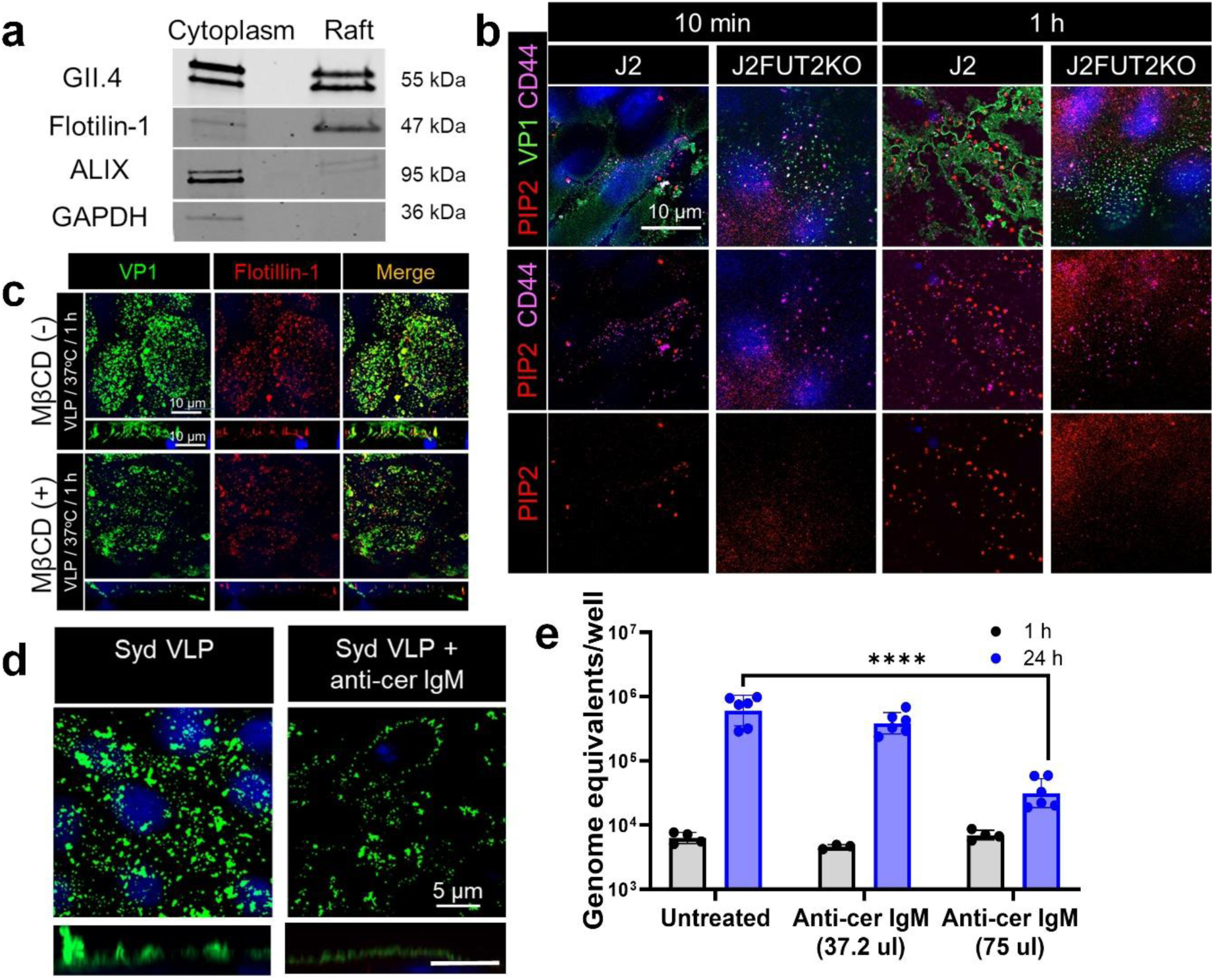
Lipid rafts and ceramide are critical determinants of GII.4 Sydney capsid clustering and viral replication. **(a)** Western blot analysis using Gp Syd-pAb of J2 HIEs incubated with GII.4 Sydney VLPs (1 h at 37°C) showing the presence of GII.4 VP1 in both cytoplasm and lipid rafts (isolated from the plasma membranes). **(b)** Incubation of HIEs with GII.4 Sydney VLPs (green) at 37°C showing an increase in PIP2 staining (red) in only secretor positive J2 HIEs over time (10 min vs 1 h) by confocal microscopy. CD44 (purple) was used as control. **(c)** GII.4 Sydney VP1 (green) clustering and colocalization with lipid raft marker flotilin-1 (red) is reduced in the presence of cholesterol sequestrant methyl beta cyclodextran (MβCD). **(d)** Inhibition of surface exposed ceramides using mouse anti-ceramide IgM reduces GII.4 VLP-induced capsid clustering (green) observed post incubation at 37°C using Gp Syd-pAb. **(e)** Inhibition of surface ceramides by incubating J2 HIEs with mouse anti-ceramide IgM (1 h before, during and after infection) reduces GII.4 Sydney viral replication at 24 h. Error bars represent mean ± SD with significance (P values) calculated using one-way ANOVA, Dunnett’s multiple comparisons test.

We next examined the role of cholesterol in GII.4 clustering, given that it is essential for maintaining raft architecture and CLIC-mediated cargo internalization (Goswami et al., 2008). Disruption of membrane cholesterol with methyl-β-cyclodextrin (MβCD) significantly impaired VLP-induced cluster formation and abrogated VP1 co-localization with flotillin-1 (Fig. 3c). Similarly, blockade of ceramides—another class of lipid raft constituents and CLIC regulators—using a multivalent anti-ceramide IgM also reduced VP1 clustering (Fig. 3d) and suppressed GII.4 replication (Fig. 3e), further confirming the requirement for intact lipid rafts in facilitating viral uptake.

Together, these data support a model in which GII.4 capsid clustering and entry are driven by HBGA-dependent recruitment of VP1 into cholesterol- and ceramide-rich glycan containing lipid rafts, which act as organizing platforms for CLIC-mediated endocytosis. Because protein glycosylation can modulate cargo–lectin interactions during raft-dependent, clathrin-independent uptake (Lakshminarayan et al., 2014; Mathew & Donaldson, 2018), we also examined the role of glycan processing in Sydney infection. Treatment of HIEs with the N-glycosylation inhibitor Kifunensine (Kif) or the O-glycosylation inhibitor Benzyl-α-GalNAc (Benz) had no effect on VLP clustering (Supplementary Fig. 2a), but both significantly reduced viral replication (Supplementary Fig. 2b), indicating that host protein glycosylation is dispensable for clustering but essential for productive infection. The efficacy of inhibitors in HIEs was confirmed by validating its affect on CD13, a N-and O-glycosylated protein (Supplementary Fig. 2c).

### Capsid integrity, avidity and glycan specificity are crucial for GII.4-induced membrane wounding and entry

Our previous work established that initial attachment of GII.4 Sydney VLP with fucosylated HBGAs triggers membrane wounding and downstream signaling (Ayyar et al., 2023). In contrast, secretor-dependent or independent, bile acid–sensitive strains such as GI.1 and GII.3 VLPs, as well as isolated GII.4 P-domain, essentially failed to induce membrane wounding or clustering (Fig. 4a; Supplementary Fig. 3a), underscoring a unique specificity of GII.4 capsid–mediated entry.

**Figure 4.**
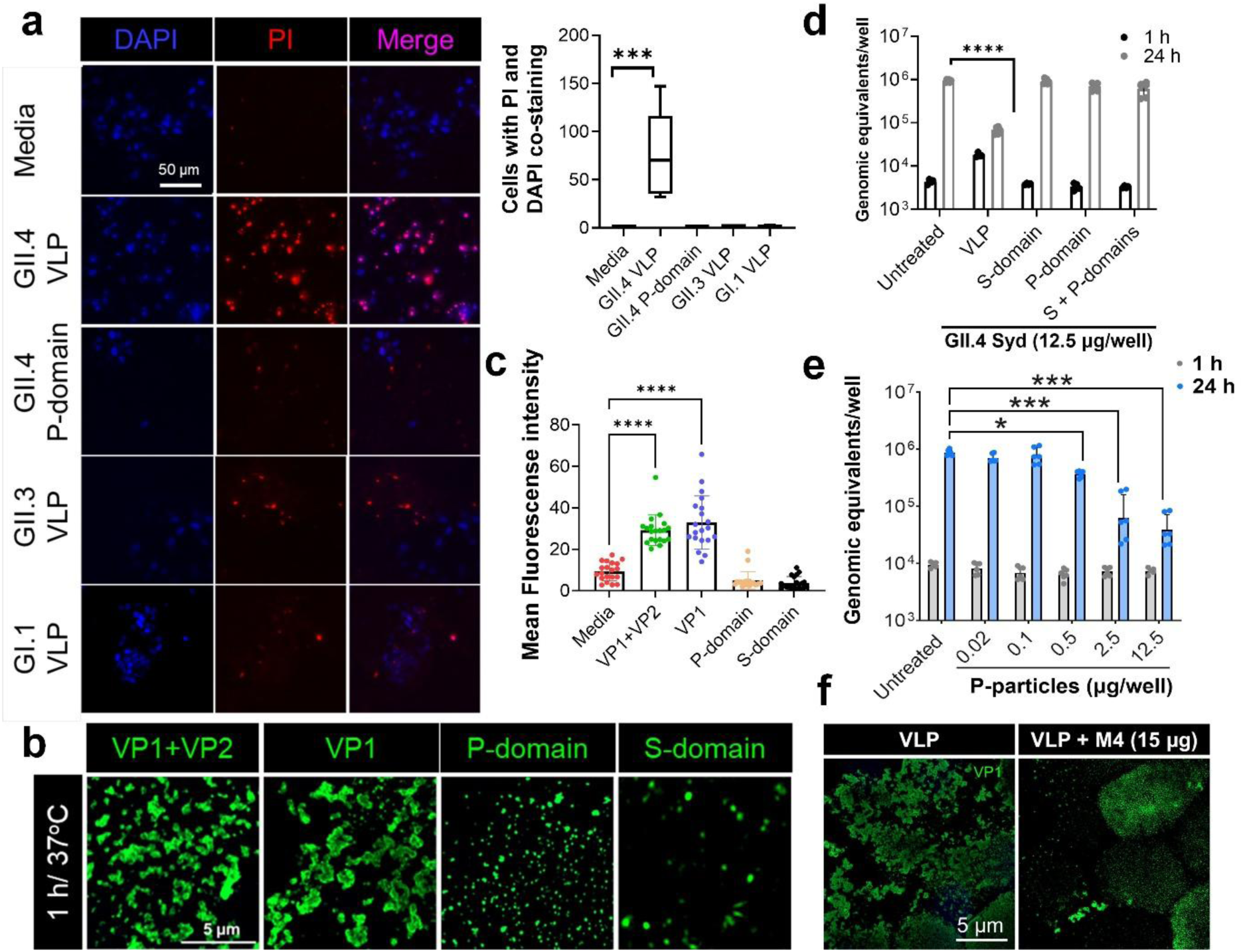
Only intact GII.4 VLPs with flexible P domain conformation can trigger cell wounding and orchestrate the capsid clustering essential for successful cell entry. (**a)** Propidium iodide (PI) uptake assay showing cell injury (post 10 min VLP incubation at 37°C) using epifluorescence microscopy. Right panel: Box and whiskers plot showing membrane injury quantitation when HIEs were incubated with media, GII.4 VLPs, GII.4 P-particles, GII.3 VLPs and GII.3 VLPs (ROI=4-6). Significance was calculated using one-way ANOVA, Dunnett’s multiple comparisons test. **(b)** Confocal microscopy showing that cluster formation in HIEs is induced by intact VP1+VP2 and VP1 only VLPs and not by shell (S-) or protruding (P) domains. **(c)** Quantitation of FM-1-43X uptake showing endocytosis is induced by VP1+VP2 and VP1 only VLPs (>1 log) and not by GII.4 S-or P-domains after 10 min at 37°C. **(d)** RT-PCR quantification of GII.4 Sydney replication (at 24 h) showing that replication is significantly inhibited in the presence of intact VLPs (VP1+VP2) and not by the presence of individual S- and P-domains or their combination. Error bars represent mean ± SD with P values calculated using one-way ANOVA, Dunnett’s multiple comparisons test for 24 h (n=3). **(e)** Inhibition of GII.4 Sydney viral replication in J2 HIEs in the presence of P-particles quantified by RT-PCR using two biological replicates for 1 h and three biological replicates for 24 h with two technical replicates for each condition (n=2). P values were calculated using Unpaired T-test for 24 h representing viral replication. *P ≤ 0.05, ***P ≤ 0.001. **(f)** Confocal microscopy showing inhibition of GII.4 Sydney-induced cluster formation by a non-HBGA blocking M4 nanobody that binds capsids in a raised conformation.

To dissect the contribution of individual capsid components of the GII.4 Sydney, we tested various constructs including full-length capsid VP1+VP2 VLPs, VP1-only particles (Table 2), isolated P- and shell (S-) domains, and high-avidity recombinant P-particles (Tan & Jiang, 2005) (Supplementary Fig 3b, 3c). Although HBGA binding is mediated by the P-domain (Cao et al., 2007; Shanker et al., 2011), only intact GII.4 VLPs triggered significant membrane wounding, clustering and endocytosis (Fig. 4a; 4b and 4c). Neither GII.4 P-particles nor isolated P- or S-dimers induced these responses, despite their ability to bind HBGAs (Fig. 4c, Supplementary Fig. 3c-3f) or cellular host factors (Ayyar et al., 2023).

**Table 2:** List of VLPs used in the study.

| **Table 2: List of VLPs used in the study** |  |  |  |  |
| --- | --- | --- | --- | --- |
| **Genotype** | **Variant** | **Year** | **GenBank**<br>**Accession No.** | **VLP**<br>**contains** |
| **GII.4** | Grimsby (GRV) | 1995 | AJ004864 | VP1 |
| **GII.4** | Lanzou (HoV) | 2002 | EU310927 | VP1+VP2 |
| **GII.4** | Farmington Hills<br>(FH) | 2002 | AY502023 | VP1+VP2 |
| **GII.4** | Farmington Hills<br>(FH-MD) | 2004 | DQ658413 | VP1 |
| **GII.4** | Hunter | 2004 | AB303941 | VP1 |
| **GII.4** | Sakai | 2005 | AB220922 | VP1 |
| **GII.4** | Yerseke | 2006 | EF126963 | VP1 |
| **GII.4** | Den Haag | 2006 | EF126965 | VP1 |
| **GII.4** | Apeldoorn (AP) | 2008 | AB541274 | VP1 |
| **GII.4** | New Orleans (NO) | 2009 | GU445325 | VP1 |
| **GII.4** | Sydney | 2012 | JX459908 | VP1+VP2 |
| **GII.4** | Sydney VP1 | 2012 | JX459908 | VP1 |
| **GII.4** | Sydney V333A<br>mutant | 2012 | NA | VP1+VP2 |
| **GII.4** | Sydney R339A<br>mutant | 2012 | NA | VP1+VP2 |
| **GII.4** | Sydney R345A<br>mutant | 2012 | NA | VP1+VP2 |
| **GII.3** | TCH04-577 | 2004 | AB365435 | VP1+VP2 |
| **GI.1** | Norwalk | 1968 | M87661 | VP1+VP2 |

In addition, clustering and uptake require only the major VP1 capsid protein, with VP2 having no measurable effect (Fig. 4b, 4c). Intriguingly, while P-particles lacked entry competence (Supplementary Fig. 3e), they inhibited viral replication in HIEs, unlike the P-domain dimer (Fig. 4d, 4e), suggesting that glycan-binding avidity alone is insufficient for entry without proper capsid architecture.

Supporting a role for capsid integrity and flexibility in virus entry, pre-treatment with the M4 nanobody—characterized as recognizing an epitope distal to the HBGA binding site, accessible only in the raised capsid conformation that triggers particle disassembly (Salmen et al., 2023)—disrupted clustering and entry into HIEs (Fig. 4f). Together, these findings emphasize that GII.4 VLP-induced clustering and endocytosis are driven by an intact capsid geometry with flexible P-domains, which enables high-avidity interactions and membrane remodeling critical for productive infection.

### GII.4 variants exhibit differential capsid clustering, endocytosis efficiency, and entry characteristics

Building on our findings that the intact GII.4 Sydney capsid alone triggers lysosomal exocytosis and membrane wounding, which are crucial for initiating endocytosis (Ayyar et al., 2023), we next investigated whether variability exists in entry mechanisms between bile acid-dependent and - independent HuNoV genotypes, and within different GII.4 variants. Pre-incubation of HIEs with an anti-LAMP-1 antibody confirmed LAMP-1’s critical role in both GII.4 endocytosis (Fig. 5a) and infection (Fig. 5b). In contrast to bile acid-dependent GII.3 and other caliciviruses (Murakami et al., 2020; Shivanna et al., 2015), we confirmed that GII.4 Sydney VLPs alone induce both lysosomal exocytosis (Supplementary Fig. 4a) and clustering (Supplementary Fig. 3a) suggesting a cell entry feature unique to GII.4 variants. In addition, all six GII.4 variants tested were able to infect HIEs in the absence of bile acids, further supporting their bile acid-independent entry mechanism (Supplementary Fig. 4b).

**Figure 5.**
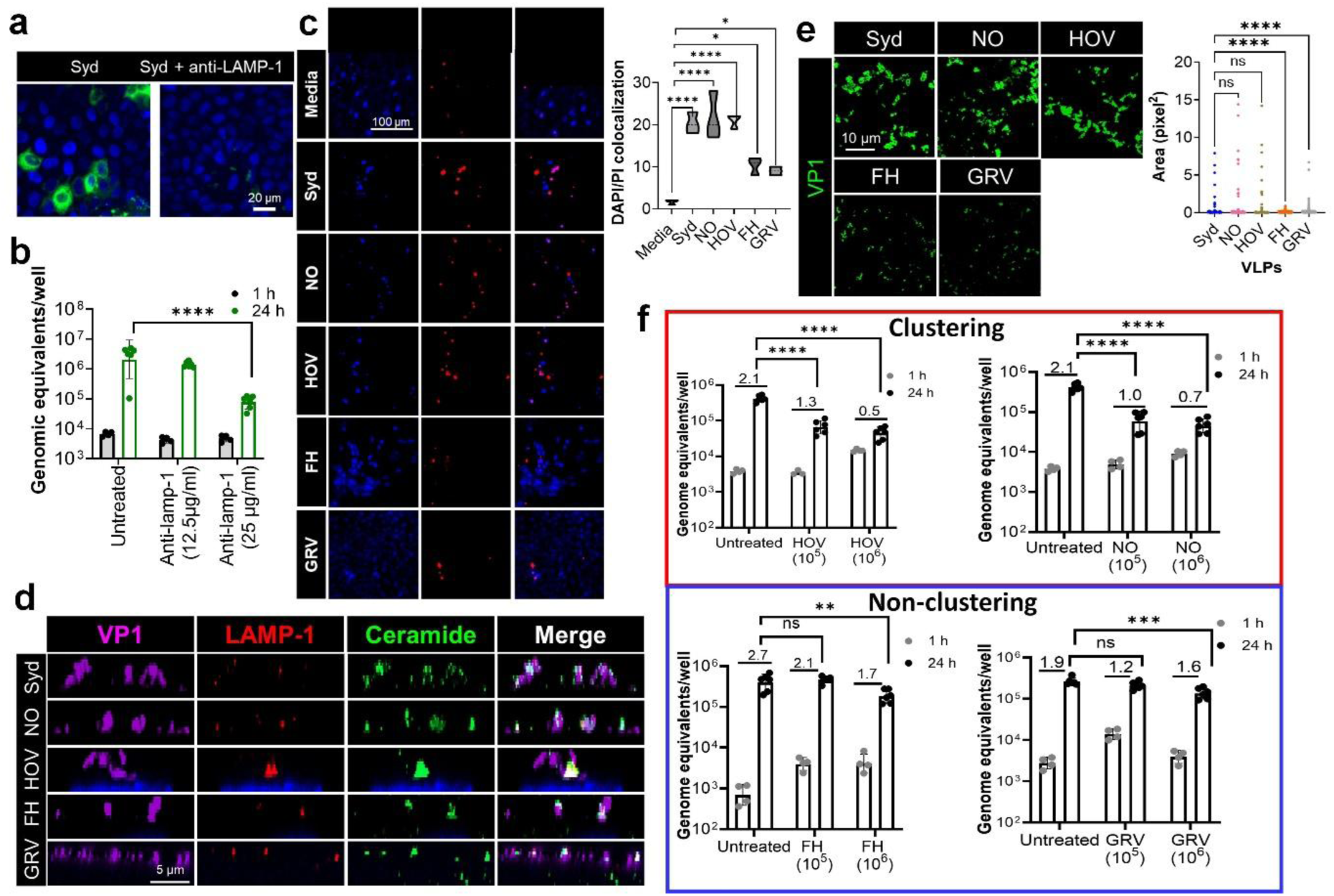
Multiple GII.4 variants induce lysosomal exocytosis but differ in clustering and membrane wounding. **(a)** Pretreatment with mouse anti-LAMP-1 mAb inhibits GII.4 Sydney-induced endocytosis (FM-1-43FX uptake, green) in J2 HIEs. **(b)** GII.4 Sydney replication is inhibited by mouse anti-LAMP-1 antibody. Quantification of replication at baseline 1 h after inoculation (n=2 HIE replicates) and at 24 h (n=3 HIE replicates) by RT-PCR. **(c)** VLP-induced wounding by different GII.4 strains using PI assay (at 10 min at 37°C). Right panel graph: Quantitation of PI/DAPI colocalized spots in 3 HIE replicates treated with media and VLPs. **(d)** VLP-induced lysosomal exocytosis in J2 HIEs 1 h post incubation at 37°C by multiple GII.4 strains [Sydney/2012 strain (SYD), Houston/2002 strain (HOV), New Orleans/2009 strain (NO), Farmington Hills/2002 (FH) and Grimsby/1995 (GRV)]. GII.4 capsid (purple), LAMP-1 (red) and ceramide (green) were stained using Gp Syd-pAb and mouse anti-LAMP-1 mAb and visualized by confocal microscopy. **(e)** Capsid-induced cluster formation (green) by a subset of different GII.4 strains in J2 HIEs detected by confocal microscopy. Right panel graph: Quantitation of cluster formation induced by GII.4 VLPs. **(f)** Competition studies evaluating GII.4 Sydney replication in the presence of VLPs from clustering (red box, HOV and NO) and non-clustering (blue box, FH and GRV) GII.4 strains. Replication inhibition was quantified at 1 h (n=2 HIE replicates) and 24 h (n=3 HIE replicates). P values calculated using one-way ANOVA, Dunnett’s multiple comparisons test (n=3).

We then extended our investigation to include VLPs of multiple GII.4 strains, such as New Orleans/2009 (NO), Houston Virus/2002 (HOV), Farmington Hills/2002 (FH), and Grimsby/1995 (GRV), to examine their entry mechanisms after validating their capsid integrity by negative stain microscopy (Supplementary Fig. 4c). Like GII.4 Sydney, all these strains induced membrane wounding (Fig. 5c), recruited LAMP-1 (Fig. 5d), and promoted exposure of ceramides at the cell surface, confirming their reliance on lysosomal exocytosis for entry. However, only a subset of the GII.4 variants (Sydney, NO, and HOV) induced capsid clustering, while FH and GRV did not (Fig. 5e). Indeed, clustering GII.4 variants exhibited enhanced endocytosis (Supplementary Fig. 4d) and stronger HBGA binding (Supplementary Fig. 4e) compared to non-clustering variants, even though all strains shared conserved amino acid in HBGA binding sites (Supplementary Fig. 4f). Infection inhibition assays further showed that VLPs from clustering strains more effectively blocked GII.4 Sydney infection (1.4 to 1.6 log_10_ reduction at 10^6^ VLP concentrations) than non-clustering strains (0.3-1 log_10_ reduction at 10^6^ VLP concentrations) (Fig. 5f), highlighting the functional importance of clustering in the entry process.

Additional testing of VP1-only VLPs from six other GII.4 variants, Farmington Hills (FH-MD; 2004), Hunter (2004), Sakai (2005), Yerseke (2006), Den Haag (2006), and Apeldoorn (2008) for membrane wounding, endocytosis, and clustering revealed that while all GII.4 variants induced wounding and endocytosis (Supplementary Figs. 5a and 5b), reduced wounding and endocytosis were observed for FH-MD and Sakai, which also exhibited less or no clustering (majority of clusters >3-5 pixel^2^), respectively (Supplementary Fig. 5a-5c). This observation led us to hypothesize that the ability to cluster might correlate with the differences in endocytic efficiency and receptor binding. These findings suggest that GII.4 variants can be categorized into clustering and non-clustering groups, and that this classification reflects differences in HBGA binding characteristics, endocytic efficiency, and overall cell entry mechanisms.

### Neighboring residues near the HBGA binding region influence GII.4-induced clustering and entry

To further dissect the molecular basis for clustering versus non-clustering phenotypes, we aligned the P-domain sequences of multiple GII.4 strains—including recent San Francisco and Wichita isolates—and observed a striking correlation at residue 333. In contrast to clustering strains encoding V333, the non-clustering strains encode M333 (Fig. 6a). Further, positions 339 and 345 remained positively charged (Arg or Lys), with R345 already known to be essential for HBGA binding based on HBGA-capsid structures (Tan et al., 2009). We hypothesized that V333 and R339 might contribute to membrane destabilization and wounding.

**Figure 6.**
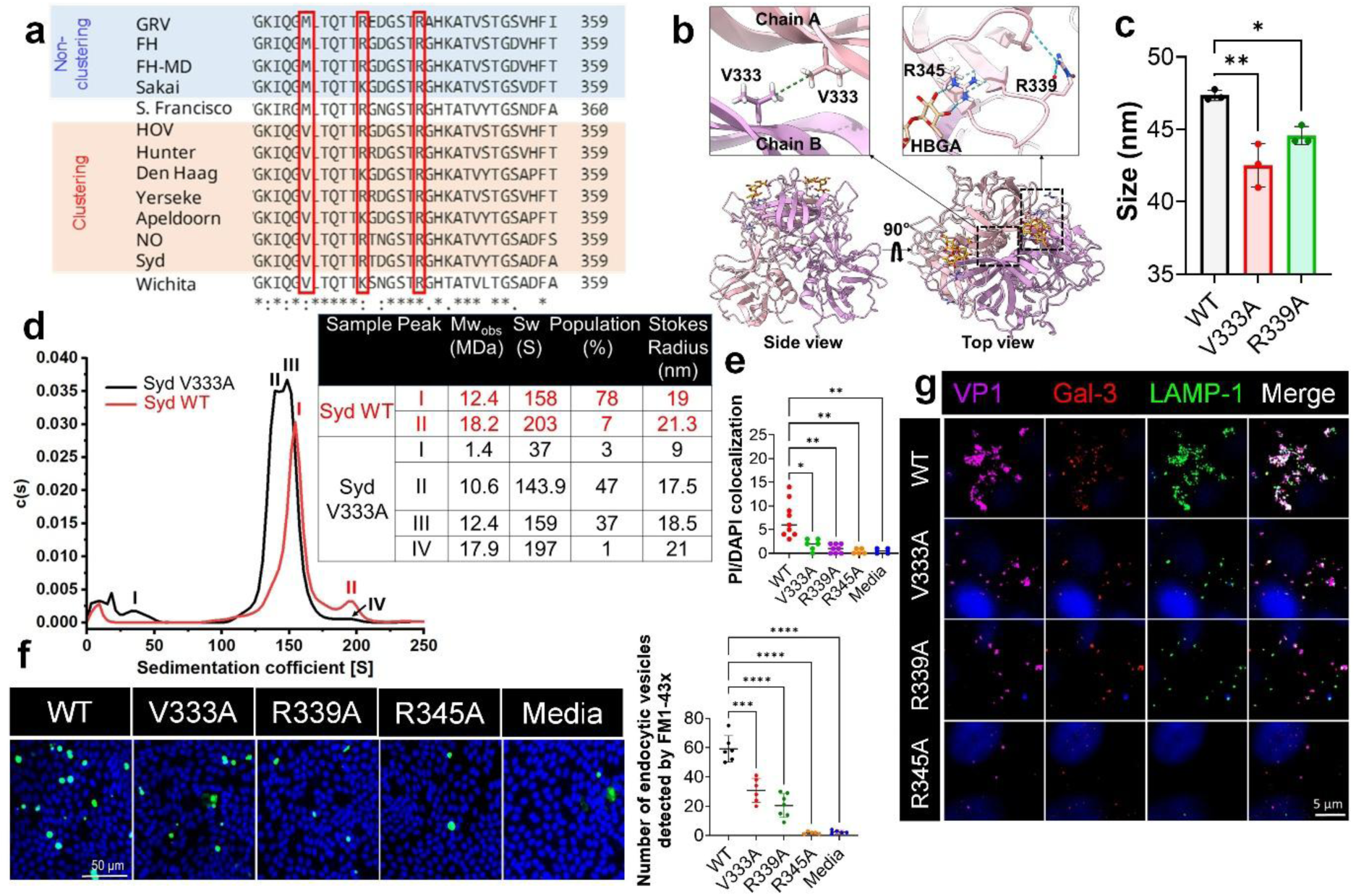
Structural and functional heterogeneity among GII.4 VLP mutants affects cell entry. **(a)** Multiple sequence alignment of sequences from clustering and non-clustering GII.4 strains using ClustalW. The amino acids targeted for preparing mutants are boxed in red. **(b)** Illustration of the wild-type (WT) GII.4 Sydney VP1 P dimer bound to fucose of HBGA type A (PDB: 4WZT; VP1= pink and violet, HBGA= orange, H-bonds= dashed blue lines, van der Waals contacts= dashed green line) highlighting the positions of V333, R339 and R345 residues. PDB: 4WZT; VP1 P domains: pink and violet, HBGA: orange, water: red spheres, H-bonds: dashed blue lines, van der Waals contacts: dashed green line. **(c)** Dynamic light scattering (DLS) measurement of the hydrodynamic diameter of WT and specified GII.4 Sydney mutant VLPs in PBS at pH 6. **(d)** Sedimentation velocity-analytical ultracentrifugation analysis (SV-AUC) of WT (red) and mutant V333A VLP (black), performed using a Beckman-Coulter XL-A analytical ultracentrifuge. The table summarizes the different parameters obtained from SV-AUC analysis. **(e)** Quantitation of VLP-induced wounding by WT and mutant VLPs in J2 HIEs (ROI=4-9) using the PI assay (at 10 min at 37°C). P values were calculated using one-way ANOVA, Dunnett’s multiple comparisons test (n=3). **(f)** Endocytosis (FM-1-43X uptake) elicited by WT, V333A, R339A and R345A VLPs post 10 min VLP treatment. (ROI=5-7). Right side: quantitation of FM-1-43X positive compartments induced by GII.4 VLPs (WT and mutants). Error bars represent mean ± SD with P values calculated for using one-way ANOVA, Dunnett’s multiple comparisons test (n=3). (g) Capsid-induced cluster formation (purple) and colocalization of VP1 with Gal-3 (red) and LAMP-1 (green) in J2 HIEs inoculated with GII.4 Sydney WT and mutant VLPs.

To test the role of specific VP1 residues in norovirus clustering and entry, we generated three GII.4 Sydney VLP mutants: V333A and R339A, targeting putative post-attachment entry regulators, and R345A as an HBGA non-binding control (Fig. 6a, b). All mutants successfully assembled into VLPs, comparable in morphology to WT VLPs, as confirmed by electron microscopy (Supplementary Fig. 6a). HBGA-binding assays showed that both V333A and R339A retained glycan-binding activity, while R345A lost binding capacity, consistent with R345’s established role in hydrogen bonding with the fucose moiety of HBGAs (Supplementary Fig. 6b).

Dynamic light scattering (DLS) analysis revealed monodisperse profiles across all samples. WT VLPs exhibited an average hydrodynamic diameter of ∼45 nm, consistent with a mixed T=3/T=4 architecture seen in cryo-EM. In contrast, V333A and R339A mutants displayed slightly reduced diameters (∼41 nm), which may reflect altered T=3/T=4 capsid ratios, a conformational shift between resting and raised states, or both (Fig. 6c). V333 is positioned at the P-dimer interface and, unlike M333 in the FH strain, engages in hydrophobic interactions with the corresponding V333 from the neighboring monomer—possibly explaining the compaction observed in the V333A mutant (Fig. 6b).

Sedimentation velocity analytical ultracentrifugation (SV-AUC) further characterized these structural changes. WT VLPs migrated predominantly (78%) as a single species with a Stokes radius of 19 nm. In contrast, V333A VLPs exhibited two distinct peaks accounting for 47% and 37% of the population, with radii of 17.5 and 18.5 nm, respectively, indicating a significant reduction in molecular weight and potential dimer destabilization (Fig. 6d). R339A VLPs aggregated during SV-AUC and could not be analyzed by this method.

Functionally, both V333A and R339A mutants exhibited markedly reduced membrane wounding and endocytic uptake compared to WT, while R345A was entirely inactive (Fig. 6e, f). None of the mutants triggered VP1 clustering at the apical membrane or colocalized with Gal-3 or LAMP-1 (Fig. 6f, g), supporting their defect in downstream entry processes.

R339 resides in a flexible surface loop near the HBGA-binding pocket and may participate in interactions with membrane components or co-receptors—potentially explaining the functional loss when mutated. In contrast, the mechanism underlying V333A’s defect is less direct. Comparative structural analysis with murine norovirus (MNV) reveals that mutation of the analogous residue V339 to isoleucine stabilizes a resting conformation of the capsid in the absence of bile salts or metal ions. Although HuNoV V333 and MNV V339 occupy the same β-strand, they are not structurally equivalent: MNV V339 points away from the P-dimer interface (Sherman et al., 2025). In silico hydrophilicity mapping shows that substituting Sydney V333 with alanine or methionine increases local hydrophilicity at the P-dimer interface (Supplementary Fig. 6c), potentially weakening van der Waals interactions and destabilizing dimer integrity.

Together, these findings identify V333 and R339—alongside the HBGA-binding residue R345—as critical molecular determinants of GII.4 capsid clustering, membrane remodeling, and CLIC-mediated entry.

## Discussion

Receptor clustering enhances the selectivity and avidity of multivalent ligands, facilitating processes like immune signaling and cell communication (Xie et al., 2025). In viral infections, host glycans can be critical modulators of susceptibility, mediating initial attachment and tissue tropism (Sieben et al., 2020). In our study, we observed that capsid clustering by GII.4 Sydney HuNoV occurs specifically in secretor-positive intestinal cultures that support replication (Fig. 1d; Supplementary Fig. 1a–c). This clustering is initiated by HBGA binding (Fig. 1e, f), followed by recruitment of Gal-3 and LAMP-1—both essential for successful infection (Fig. 2f, g; (Ayyar et al., 2023)). These findings outline a sequential cascade in which initial glycan-dependent attachment triggers capsid reorganization, priming the apical cell membrane for virus entry.

Our results align with prior studies showing that viral clustering enhances infectivity by promoting receptor engagement, increasing local capsid concentration, and facilitating membrane remodeling (Coyne & Bergelson, 2006; New et al., 2021; Nieto-Garai et al., 2021; Wang et al., 2024; Zhang et al., 2023). Notably, GII.4 clustering is linked to glycan-induced lipid raft rearrangement (Fig. 3b), resembling mechanisms exploited by influenza A to optimize multivalent receptor binding (Sieben et al., 2020). Disruption of lipid dynamics impaired both clustering and replication (Fig. 3c–e), highlighting the importance of optimal membrane architecture in GII.4 entry. These observations echo earlier findings with HBGA-conjugated giant unilamellar vesicles (GUVs), where GII.4 VLP engagement alone was sufficient to drive glycosphingolipids toward the interaction site (Rydell et al., 2009), thus strenghthening viral-host binding. This interaction ultimately promoted capsid clustering and membrane deformation, independent of cellular machinery (Parveen et al., 2018; Rydell et al., 2013). Similar lipid-driven clustering has been reported for diverse viruses such as SV40, influenza and MNV (Dahmani et al., 2019; Kociurzynski et al., 2019; Stewart et al., 2025), reinforcing the central role of membrane composition in viral uptake.

Importantly, the structural multivalency of VLPs—comprising 180 VP1 subunits arranged with precise capsid geometry as 90 dimers—confers high glycan avidity and functionally distinguishes them from recombinant P-domains or P-particles. Although both retain HBGA-binding capability, only intact VLPs are capable of inducing membrane wounding and nanoscale clustering (Fig. 4a, 4b; Supplementary Fig. 3d, 3f), highlighting that capsid integrity and overall architecture with increased avidity are critical for engaging host membranes and initiating entry. Structural studies have shown that the HuNoV capsid is dynamic, with flexible hinges at the P-domain facilitating conformational shifts that may regulate receptor engagement and downstream entry events (Hu et al., 2022), similar to mechanisms first described in MNV (Sherman et al., 2019). Notably, conformational flexibility induced by glycan binding is crucial for subsequent receptor interactions, as observed with HPV16, where initial heparan sulfate binding primes the virus for engagement with secondary receptors (Feng et al., 2024). Thus, in GII.4 HuNoVs, capsid plasticity likely enables spatial reorganization of host factors and membrane remodeling, underscoring the central role of structural dynamics in promoting efficient viral entry.

All these features are also true with GII.3, yet they do not cluster, supporting a model that GII.4 HuNoVs have evolved a uniquely potent entry mechanism (Ayyar et al., 2023). Unlike bile acid-dependent norovirus genotypes such as GII.3, GI.1 or GII.17, GII.4 Sydney VLPs trigger robust membrane wounding and capsid clustering (Fig. 4a, Supplementary Fig. 3a) facilitating viral entry and replication. This functional divergence may reflect distinct virus biology, co-receptors, entry routes and host immune signatures, consistent with recent studies showing that the endocytic pathway modulates innate immune responses (Abad & Danthi, 2022) and contributes to genotype-specific infection outcomes (Lin et al., 2020; Murakami et al., 2020; Prasad et al., 2025).

To explore the conservation of these features, we assessed entry characteristics of multiple GII.4 variants. Variants including HOV, NO, Apeldoorn, Yerseke and Den Haag, induce pronounced apical wounding and assemble into dense VP1 clusters, whereas, earlier chronologically appearing strains such as FH and GRV show markedly reduced wounding and clustering (Fig 5c, e; Supp Fig 5a, c). Quantitative analysis revealed a strong correlation between the degree of clustering, endocytic uptake, and HBGA-binding (Supplementary Fig. 4c, d, 5b), resembling how GM1 clustering enhances membrane deformation and internalization of cholera toxin (Kabbani et al., 2020; Wolf et al., 2008). These findings align with recent observations that membrane disruption facilitates viral infectivity (Jiao et al., 2024) and suggest that ligand-induced nanodomain formation primes the apical surface for CLIC-mediated internalization.

Targeted mutagenesis of the VP1 P domain—V333A or R339A—uncouples glycan binding from entry: both mutants bind HBGAs normally but exhibit impaired membrane remodeling and endocytic uptake (Fig. 6a; Supplementary Fig. 6a-c). This shows that after initial attachment, precise capsid geometry and flexibility are essential to engage downstream host factors—presumably an as-yet-unidentified co-receptor.

This multi-step entry strategy utilized by GII.4 parallels other non-enveloped viruses that exploit receptor-or environment-triggered capsid rearrangements to breach host membranes and deliver their genomes. For example, poliovirus and adenovirus expose internal hydrophobic loops upon receptor binding to form membrane-permeabilizing pores (Fricks & Hogle, 1990; Wiethoff & Nemerow, 2015); rotavirus and bluetongue virus undergo proteolysis-induced conformational shifts that drive membrane penetration (Dormitzer et al., 2004; Xia et al., 2021); and rhinovirus VP4 externalizes to assemble size-selective channels (Panjwani et al., 2014), while both reovirus and MNV norovirus exploit cholesterol-dependent, lipid-centric pathways for cellular uptake (Snyder & Danthi, 2016; Stewart et al., 2025). Together, these examples define a conserved paradigm in which discrete capsid rearrangements synergize with lipid remodeling to mechanically and biochemically subvert host plasma membrane.

In this context, GII.4 noroviruses orchestrate a multistep internalization program by coupling high-avidity HBGA binding with cholesterol- and ceramides-driven raft remodeling and VP1-mediated nanoscale clustering. Our delineation of clustering versus non-clustering phenotypes—centered on residues V333 and R339—pinpoints the molecular determinants underpinning the GII.4 Sydney infectivity profile. While V333 and R339 are critical for GII.4 Sydney’s entry efficiency, other viral factors—such as mutations that alter ligand-binding preferences, antigenicity or domain chemistry/conformation, and changes in the polymerase—likely collaborate to shape infectivity, fitness, and pandemic potential of the virus (Barclay et al., 2025; Tohma et al., 2024). Although further analyses are required to define how these residues govern capsid–host interactions, our findings deepen the mechanistic understanding of HuNoV entry and prioritize discrete viral and host factors for future co-receptor discovery and targeted therapeutic development.

## Methods

### Virus and VLPs

The virus used in these studies was TCH12-580, a GII/Hu/US/2012/GII.4 Syd [P31]/TCH12–580 strain. Stool filtrates were prepared as 10% suspensions in phosphate-buffered saline (PBS) with viral titer determined by real-time RT-PCR as previously described (Ettayebi et al., 2016; Ettayebi et al., 2024). VLPs were produced in a baculovirus expression system using open reading frame 2 (ORF2) + ORF3+ untranslated region (UTR) sequences for different HuNoV strains (Table 2). VP1-only VLPs Farmington Hills (FH-MD; 2004), Hunter (2004), Sakai (2005), Yerseke (2006), Den Haag (2006), and Apeldoorn (2008) were generously provided by Dr. Gabriel Parra from the Division of Viral Products, Food and Drug Administration, USA (Kendra et al., 2021).

Mutant VLPs were generated by co-transfecting 0.9 x10^6^ Sf9 insect cells with the pVL1393 transfer vector, harboring ORF2+ORF3+3’UTR of a GII.4 Sydney, and the linearized BestBac™ baculovirus DNA (#91-200, Expression Systems). Transfected cells were incubated for 5 days to generate p0 baculovirus stocks. For recombinant protein expression, p0 virus was amplified to make larger baculovirus stocks (titer 3.0-3.2 x 10^8^ PFU/mL) that were used to infect Sf9 insect cells with a MOI 5 for 10 days. Intact VLPs were purified as previously described using CsCl gradient centrifugation (Jiang et al., 1992).

### Preparation of HIE monolayers

HIE cultures used in the current study were described previously with recent optimization (Ettayebi et al., 2016; Ettayebi et al., 2024). Most studies used the permissive, secretor positive jejunal (J2) HIE line. An isogenic, genetically modified J2FUT2KO HIE that lacks FUT2 expression as previously described (Haga et al., 2020) was used for comparison with J2 HIEs. A list of all HIE lines used in this study is provided in Table 1.

To prepare cultures for infections, 3D HIE cultures were dissociated by trypsinization followed by gentle pipetting and passing the cells through a 40 μm cell strainer. Cells were pelleted and resuspended in OGM proliferation media (#06010, Stem Cell Technologies) supplemented with 10 μM ROCK inhibitor Y-27632 and plated as monolayers. After 24 h, OGM differentiation medium was added to the monolayers and the cells were allowed to differentiate for 5 days with intermittent media change (Ettayebi et al., 2024).

### VLP labelling

VLPs were labeled using Alexa Fluor™ 488 Protein Labeling Kit (#A10235, ThermoFisher Scientific) as per manufacturer’s instructions. Briefly, 0.5 mL of the 2 mg/mL protein solution was mixed with 50 µl of 1 M sodium bicarbonate and mixed with reactive dye mixture for 1 h at room temperature followed by purification using spin columns provided with the kit. Labelled protein was aliquoted and stored at - 20°C.

### VLP binding using confocal microscopy

HIE monolayers were grown in 8-well 15 µ-slides (#80826, Ibidi) for imaging. For binding studies, five days differentiated HIEs were treated with 1 x 10^12^ particles of VLPs for 1 h at 4°C. For studying VLP interaction with HIE over time, the VLPs were incubated with HIEs at 37°C for 10 min and 1 h. The cells were fixed with 4% paraformaldehyde (PFA) for 20 min at room temperature, blocking with 5% BSA in 0.1% Triton X-100 (for permeabilization) in PBS for 30 min at room temperature. The cells were washed with PBS and imaging was done, if labelled VLPs were used. In the case of unlabeled VLPs, the cells were incubated overnight at 4°C with primary antibodies. HuNoV capsid protein (VP1) was detected using 1:1000 dilution of guinea pig anti-Sydney polyclonal Ab (Gp Syd-pAb), whereas, LAMP-1, flotillin-1 and ceramides were detected using 1:200 dilution of mouse anti-LAMP-1 (#sc-20011, Santa Cruz Biotechnology), mouse anti-flotillin-1 (# 610821, BD Biosciences) and 1:100 dilution of rabbit anti-ceramide pAb (Krishnamurthy et al., 2007). After washing (3 times, 10 min each), the cells were incubated with 1:500 dilution of donkey anti-mouse 549 (#610-742-124, Rockland), donkey anti-rabbit 649 (#611-743-127, Rockland), and anti-guinea pig 488 secondary antibodies (#606-141-129, Rockland). UEA1 staining was done using 1:500 dilution of rhodamine-labelled UEA1 (#RL-1062-2, Vector Laboratories). UEA1 inhibition assay was carried out by incubating the HIEs with unconjugated UEA1 (L-1060, Vector Laboratories) 1 h at 37°C prior to VLP treatment. The cells were washed three times, and nuclei were stained with 4, 6-diamidino-2-phenylindole (DAPI) (300 nM) for 5 min at room temperature followed by subsequent Z-stack images captured using either GE Healthcare DeltaVision Deconvolution Microscope or Zeiss Laser Scanning Microscope LSM 980.

### Time lapse microscopy

Time lapse microscopy was carried out by adding fluorescently labeled VLPs to J2 HIE monolayers and capturing subsequent Z-stack images every 2.5 mins for a period of 1 h at 37°C using a GE Healthcare DeltaVision Deconvolution Microscope. The images were converted into a movie using ImageJ AVI plugin.

### HuNoV infection of J2 HIE monolayers

HuNoV infection was performed with HIEs with/without the addition of bile acid GCDCA (500 µM) (Ettayebi et al., 2016; Ettayebi et al., 2024). Briefly, 9 × 10^5^ GEs of GII.4 stool filtrate in OGM differentiation medium was added to the HIEs and incubated for 1 h at 37°C. After incubation, the monolayers were washed twice with CMGF(-) and were incubated for 24 h in the differentiation medium. Viral replication was quantified by Reverse Transcriptase Quantitative Polymerase Chain Reaction (RT-qPCR) at 24 h using 1 h binding as reference. For infection assay with inhibitors/competitors, those were added to the monolayer 1 h prior to infection, during infection and after infection until the cells were harvested for RNA isolation except for glycosylation inhibitors which were added to the cells since their differentiation until harvesting (5 days).

### RNA extraction and RT-qPCR

Total RNA extraction from each well was done using the KingFisher Flex Purification System and MagMAX™ Pathogen RNA/DNA Kit. RT-qPCR was carried out using the primer pair COG2R /QNIF2d and probe QNIFS with qScript XLT One-Step RT-qPCR ToughMix reagent containing ROX reference dye. RT-qPCR was performed on an Applied Biosystems StepOnePlus thermocycler using the following conditions: 50°C (15 min), 95°C (5 min), followed by 40 cycles of 95°C (15 sec) and 60°C (35 sec). A standard curve based on a recombinant HuNoV HoV RNA transcript was used to quantitate viral GEs in RNA samples.

### Wounding and endocytosis assays

Differentiated HIE monolayers were incubated with VLP (20 µg) and 5 µg/ml propidium iodide (PI) for wounding assays for 10 min at 37°C. PI was substituted with FM1-43X dye for endocytosis assays. After washing (twice), the cells were fixed with 4% PFA for 20 min followed by counterstaining with DAPI for 5 min. PI/DAPI colocalization was measured to determine VLP-induced wounding. For endocytosis, FM1-43X stained organelles in DAPI stained cells were counted to measure endocytosis.

### Clustering assays

Differentiated HIEs on slides were treated with VLPs (2 ug) and incubated for 1 h at 37°C. The cells were washed and fixed with 4% PFA for 20 min followed by permeabilization with PBS containing 0.1% Triton X-100. The cells were incubated overnight at 4°C with GP-Syd-pAb followed by probing with labelled-anti-guinea pig secondary antibody. Cluster size was calculated using measure particle size feature in imageJ/FIJI software.

### Lipid raft isolation

HIE monolayers were plated in 24-well plate by seeding 5 x 10^6^ cells/ well. After 5 days of differentiation, the cells were incubated with VLP for 1 h 37°C. After washing with CMGF(-) twice, lipid rafts from VLP-treated and untreated HIE monolayers were isolated using Minute™ Total Lipid Raft Isolation Kit (#LR-039, Invent Biotechnologies Inc.) as per manufacturer’s instructions. Purity of lipid rafts were confirmed by Western blot analysis using specific markers for cytoplasmic fraction (GAPDH) and lipid raft fraction (flotillin-1).

### P particle expression

The GII.4 Sydney P domain (Norovirus Hu/GII.4/Sydney/NSW0514/2012, amino acids 225-539) was cloned into the pMal-C2E vector with a C-terminal GSDCRGDCFC linker (Fang et al., 2013) and an N-terminal tag comprising a hexahistidine-maltose binding protein-tobacco etch virus cleavage site (6xHis-MBP-TEV). The protein was expressed in *E. coli* BL21 (DE3) cells and purified with Ni-NTA resin. The MBP moiety was removed by TEV cleavage during overnight dialysis (20 mM Tris pH 7.4, 150 mM NaCl, 1% beta-mercaptoethanol) and the cleaved P domain was separated from MBP by negative affinity chromatography on Ni-NTA resin (displaced with 20 mM Tris pH 7.4, 200 mM NaCl, 30 mM imidazole, 1% beta-mercaptoethanol) and subsequently on amylose resin (New England Biolabs). Finally, P particles were separated by size exclusion chromatography on a Superdex200 column (20 mM Tris pH 7.4, 150 mM NaCl) and their molecular weight was determined.

### HBGA binding ELISA

Direct binding of VLP to HBGAs was determined by ELISA. Porcine gastric mucin (100 μg/ml)-coated and blocked plates were incubated with different VLP concentrations (25.6 µg/ml – 0.39 ng/ml) for 1.5-2 h. The plate was washed 3 times with PBST (PBS containing 0.05% Tween 20), and PBS followed by probing the interaction with Gp Syd-pAb (1:5000 dilution in 1% blocking buffer of Gp Syd-pAb for GII.4 and GII.3 VLPs, anti-Norwalk for GI.1). Horseradish peroxidase (HRP)-labeled goat anti-guinea pig secondary antibody (1:5000) was used to detect the binding using tetramethylbenzidine (TMB) substrate. Color development was quenched using phosphoric acid and absorbance was measured at 450 nm.

For binding assays, GII.4 Sydney P-dimers or P-particles were incubated with the PGM-coated (10 µg/ml) and blocked plates for 1 h at 37°C. Bound P-protein was probed with GP Syd-pAb at 1:5,000 dilution and anti-guinea pig secondary antibody conjugated to alkaline phosphatase (AP) at 1:5,000 dilution (#A5062, Millipore Sigma). Detection of AP was performed with pNPP solution (Bio-Rad) and measured by absorbance at 405 nm.

### HBGA binding Inhibition ELISA

VLP amounts capable of inhibiting VLP binding to PGM were determined by HBGA binding ELISA. Inhibition assays were carried out by incubating the half maximal effective concentration of VLPs with different concentrations of the PGM in solution (1000 μg to 0.1 μg/ml) along with one without any PGM. After incubation at 37°C for 2 h, the VLP-PGM mixture was assayed on a PGM (100 μg/ml)-coated plate and incubated at 37 °C for 1.5-2 h, and the reaction was probed with GP Syd-pAb (1:5000) followed by detection with HRP-labeled goat anti-guinea pig secondary antibody. The percentage inhibition was calculated by subtracting the absorbance obtained for each PGM dilution (A_1_) from the absorbance without PGM (A_0_) divided by the A_0_ multiplied by 100.

### Dynamic light scattering

The hydrodynamic diameter of GII.4 Sydney WT and mutant VLPs and GII.4 Sydney P particles was measured using a ZetaSizer Nano instrument (Malvern Instruments, UK). Proteins were diluted to 200 nM in PBS, pH 6 and equilibrated at 25°C for 2 minutes. Three x 12 measurements were performed with the following standard settings: Refractive Index 1.33, viscosity 0.9, temperature 25°C. The averages of each replicate were plotted with GraphPad Prism.

### Sedimentation velocity-analytical ultracentrifugation

The SV-AUC experiments of VLP GII.4 WT and VLP GII.4 V333A were performed using a Beckman-Coulter XL-A analytical ultracentrifuge with a TiAn60 eight-hole rotor and two-channel Epon centerpieces (12 mm). Both the VLPs were prepared in PBS at pH 6.0. Absorbance scans were recorded at 280 and 220 nm at every 30-second interval at 15000 rpm at 4 °C. A Continuous distribution c(s) model was used to fit multiple scans at regular intervals with SEDFIT (Schuck et al., 2002). The solvent density (ρ) and viscosity (η) were calculated from the chemical composition of different proteins by SEDNTERP (Philo, 2023).

### Negative stain and cryo-EM screening

P particles at a concentration of 1 mg/ml were applied onto a 200-mesh R 2/2 +2 nm carbon Quantifoil Cu grid and stained with 1% uranyl acetate. Grid visualization was done on a JEM1230 microscope at 80 kV and x30,000 magnification. VLPs (3 µl) in PBS pH 6 were applied onto 300-mesh R1.2/1.3 Quantifoil Cu grids for 10 s, blotted for 4 s and plunge-frozen into liquid ethane using a Vitrobot IV(FEI) instrument at a constant humidity and temperature of 18°C. The grids were screened with a Titan Krios electron microscope fitted with a Bioquantum K2 detector at 300 kV and a magnification of x64,000.

### Statistical analysis

Each experiment was independently performed at least twice using two or more HIE replicates. For infection assays, two biological HIE replicates were used at the 1 h time point and three biological replicates at the 24 h time point. RT-qPCR assays included two or more technical replicates per biological replicate. Data analyses compared untreated and treated groups at 24 h using geometric mean values.

Statistical analyses were conducted using GraphPad Prism (v9.0). Dunnett’s multiple comparisons test was used unless otherwise specified. Depending on the experimental design, either unpaired two-tailed *t*-tests or one-way ANOVA with Tukey’s post hoc test were applied. Results are reported as mean ± standard deviation (SD). Statistical significance was defined as *p* < 0.05, with specific comparisons yielding *p*-values represented as * (*p* < 0.05), ** (*p* < 0.01), *** (*p* < 0.001), and **** (*p* < 0.0001).

## Acknowledgements

This research was supported by National Institutes of Health Grant P01 AI57788, U19 AI116497, and P30 DK56338 that supports the Texas Medical Center Digestive Diseases Core Center, S10 OD028480 that supported purchasing the Zeiss Laser Scanning Microscope LSM 980 with Airyscan 2, and the Robert Welch Foundation Q1279 grant. P30 CA125123 supported the Recombinant Protein Production and Characterization Core at Baylor College of Medicine for VLP production and AUC experiments. We acknowledge the cryo-EM Core at Baylor College of Medicine for negative stain and cryo-EM imaging and the Cryo-EM Core Facility from the UTHealth Structural Biology Imaging Center at Houston. We thank Dr. Gabriel I. Parra and Joseph A. Kendra (Division of Viral Products, Food and Drug Administration, Silver Spring, MD) for generously providing us with HuNoV VLPs, and Dr. Erhard Bieberich (University of Kentucky and VAMC, Lexington, KY) for the rabbit anti-ceramide pAb.

