## Supplementary Data for "Functional Diversity in GII.4 Norovirus Entry: HBGA Binding and Capsid Clustering Dynamics"

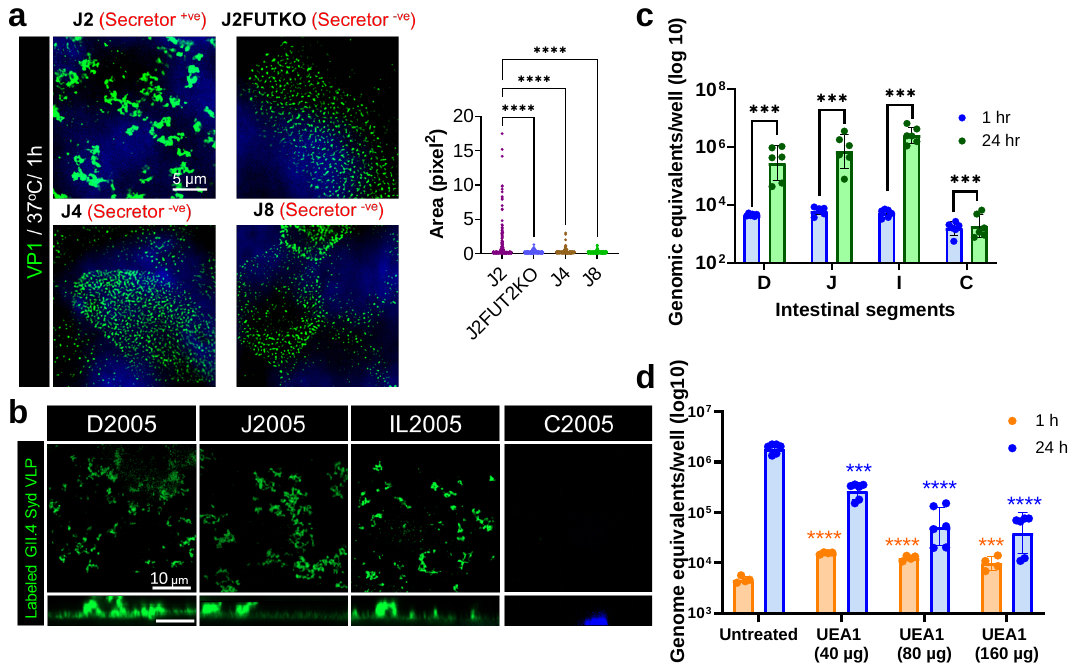

**Supplementary Figure 1. GII.4 clustering is driven by fucosylated glycans, which are tightly regulated by the expression of *FUT*2.**

**(a)** GII.4 Sydney VLP-induced clustering (green) was only observed in secretor positive (J2) HIEs and not in secretor negative HIEs (J2FUT2KO, J4 and J8) by confocal microscopy. Graph on the right represents the quantification of clusters based on area (1 pixel = 0.04 µm). **(b)** GII.4 clustering is observed in HIEs from small intestinal segments [duodenum (D), jejunum (J) and ileum (IL)] of a secretor-positive (2005) (Ettayebi et al., 2024) organ donor that also support GII.4 replication. There was minimal binding and no clustering in HIEs from the colon (C). **(c)** RT-PCR quantification of GII.4 Sydney replication in donor 2005 intestinal HIEs at 1 h and 24 h. Error bars represent mean ± SD with P values calculated using one-way ANOVA, Dunnett’s multiple comparisons test (n=3). **(d)** GII.4 Sydney viral binding and replication at 1 h (n=2 HIE replicates) and 24 h (n=3 HIE replicates) post-inoculation in the presence of UEA-1 (preincubated 1 h prior to inoculation, during inoculation and after inoculation) in J2 HIEs. Error bars represent mean ± SD with P values calculated by comparing untreated with UEA1 treated cells.

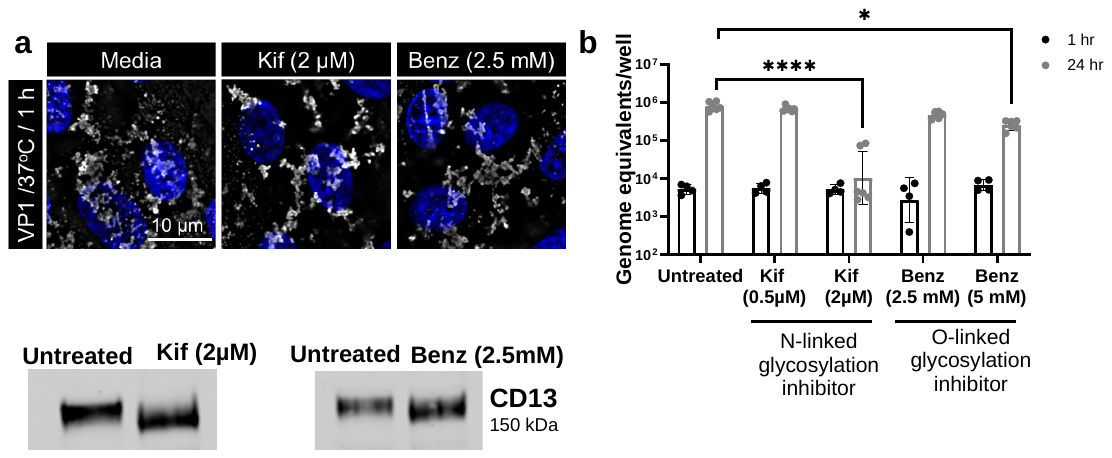

**Supplementary Figure 2. GII.4 clustering operates independently of N-linked and O-linked glycosylation, yet viral replication is dependent on N-linked and O-linked glycosylation.**

**(a)** Clustering is not inhibited in HIEs treated with N- and O-linked glycosylation inhibitors [kifunensine (Kif) and benzyl-α-GalNAc (Benz)] for four days during differentiation and infection. **(b)** GII.4 Sydney replication in the presence of N- and O-linked glycosylation inhibitors Kif and Benz, is significantly reduced. Error bars represent mean ± SD with P values calculated using one-way ANOVA, Dunnett’s multiple comparisons test (n=3). **(c)** Activity of inhibitors was confirmed by incubating the HIEs with Kif and Benz for 24 h followed by Western blotting of the cell lysates to detect N- and O-glycoprotein CD13.

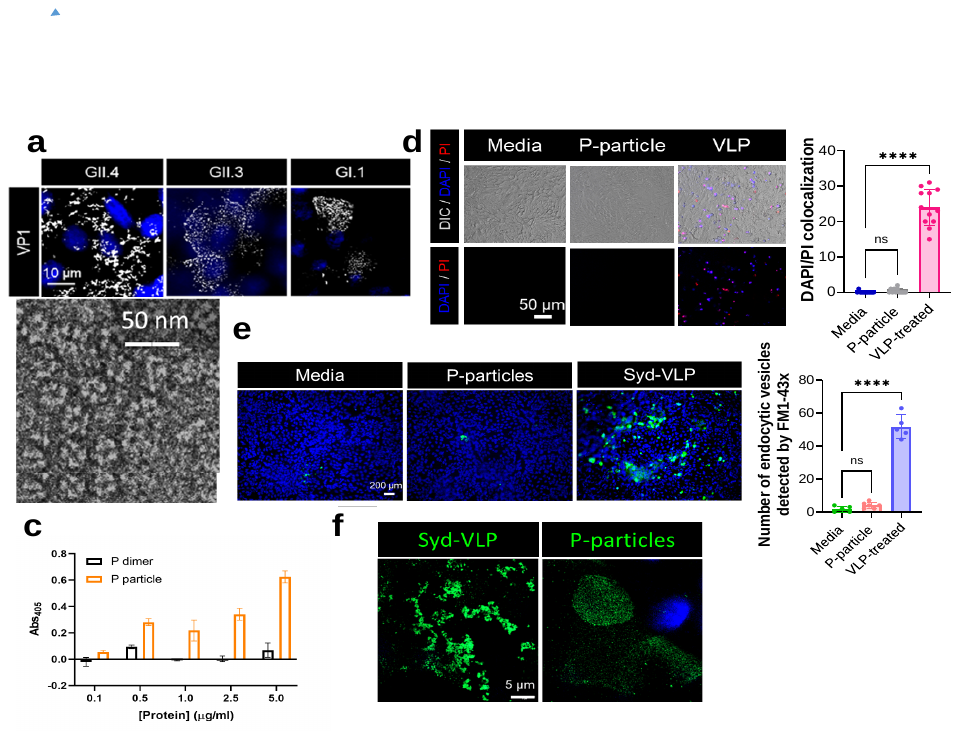

**Supplementary Figure 3. GII.4 Sydney P-particle binding to apical HBGAs is insufficient to induce membrane wounding, endocytosis or capsid clustering.**

**(a)** Confocal microscopy showing VLP clustering in J2 HIEs after the cells were incubated with VLPs of different HuNoV genotypes for 1 h at 37^o^C. **(b)** Negative stain EM image of GII.4 Sydney P-particles showing a homogeneous population of ~25 nm in diameter. **(c)** ELISA comparing the binding of GII.4 Sydney P-particles and P-dimers to HBGAs from pig gastric mucin (PGM). **(d)** Wounding of HIEs is observed only with intact GII.4 Sydney VLPs and not with P-particles. Wounding experiments were performed by incubating J2 HIEs with VLPs and P-particles in the presence of propidium iodide (PI) at 37^o^C for 10 min. Right panel: Quantification of wounding measured by counting DAPI/PI colocalization in 12 regions of interest (ROI)/condition (n=2). P values were calculated using one-way ANOVA, Sidak’s Multiple Comparisons Test, where **** P ≤ 0.0001. **(e)** Endocytosis in HIEs requires intact GII.4 VLPs. Endocytosis assays were carried out by incubating J2 HIEs for 10 min at 37^o^C with VLP and P-particles in the presence of FM1-43FX. Right panel: quantification of VLP and P-particle induced endocytosis (ROI= 5-6). P values were calculated using one-way ANOVA, Sidak’s Multiple Comparisons Test, where **** P ≤ 0.0001 (n=2). **(f)** Confocal microscopy showing intact GII.4 Sydney VLPs are required for apical clustering of viral capsids (green). Clustering was observed by incubating J2 HIEs with fluorescent VLPs and P-particles for 1 h at 37^o^C (n=2).

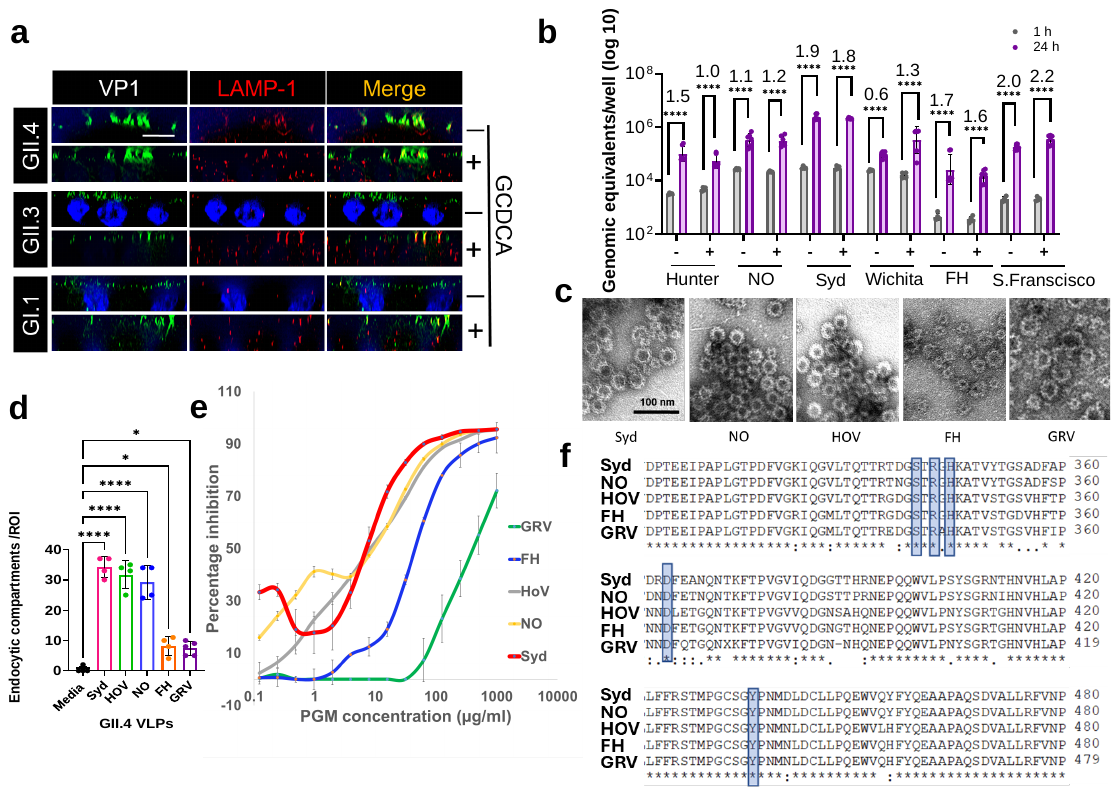

**Supplementary Figure 4. GII.4 HuNoV variants exhibit conserved HBGA binding but differential lysosomal exocytosis and endocytic efficiency.**

**(a)** VLP-induced lysosomal exocytosis in the presence and absence of bile acid GCDCA in J2 HIEs at 37^o^C (1 h) using GII.4, GII.3 and GI.1 VLPs. LAMP-1 (red) and GII.4/ GII.3 VP1 (green) colocalization (white arrows) was detected by confocal microscopy after staining with mouse anti-LAMP-1 mAb and Gp Syd-pAb. GI.1 (green) was detected using anti-GI.1 guinea pig pAb (n=3). **(b)** GII.4 replication with (+) or without GCDCA (-) in J2 HIEs. Replication was quantified by RT-PCR by comparing 1 h binding (black, n=2 replicates) to 24 h (purple, n=3 replicates) represented as fold change on the top of the bars. Error bars represent mean ± SD (**** P ≤ 0.0001) calculated using one-way ANOVA, Dunnett’s multiple comparisons test (n=3). **(c)** Negative staining images of VLPs from different GII.4 variants used in the study. **(d)** Endocytic efficiency different GII.4 variants, calculated by counting and comparing endocytic compartments (FM-1-43X uptake, green organelles) induced by their VLPs (10 min at 37^o^C), ROI=4-5. P values (**** P ≤ 0.0001, * P ≤ 0.05) were calculated using one-way ANOVA, Dunnett’s multiple comparisons test (n=3). **(e)** Avidity determination of GII.4 VLPs by inhibition ELISA. Binding of GII.4 variant VLPs to different concentrations of porcine gastric mucin (PGM) HBGAs in solution was carried out by incubating a fixed amount of VLP variant, followed by adding the mixture to the plate coated with PGM. The percentage inhibition of VLP binding to the coated HBGA by the HBGA in solution was calculated for avidity determination. **(f)** Sequence comparison showing that amino acids important for HBGA binding (represented in boxes) are conserved among the different GII.4 VLPs.

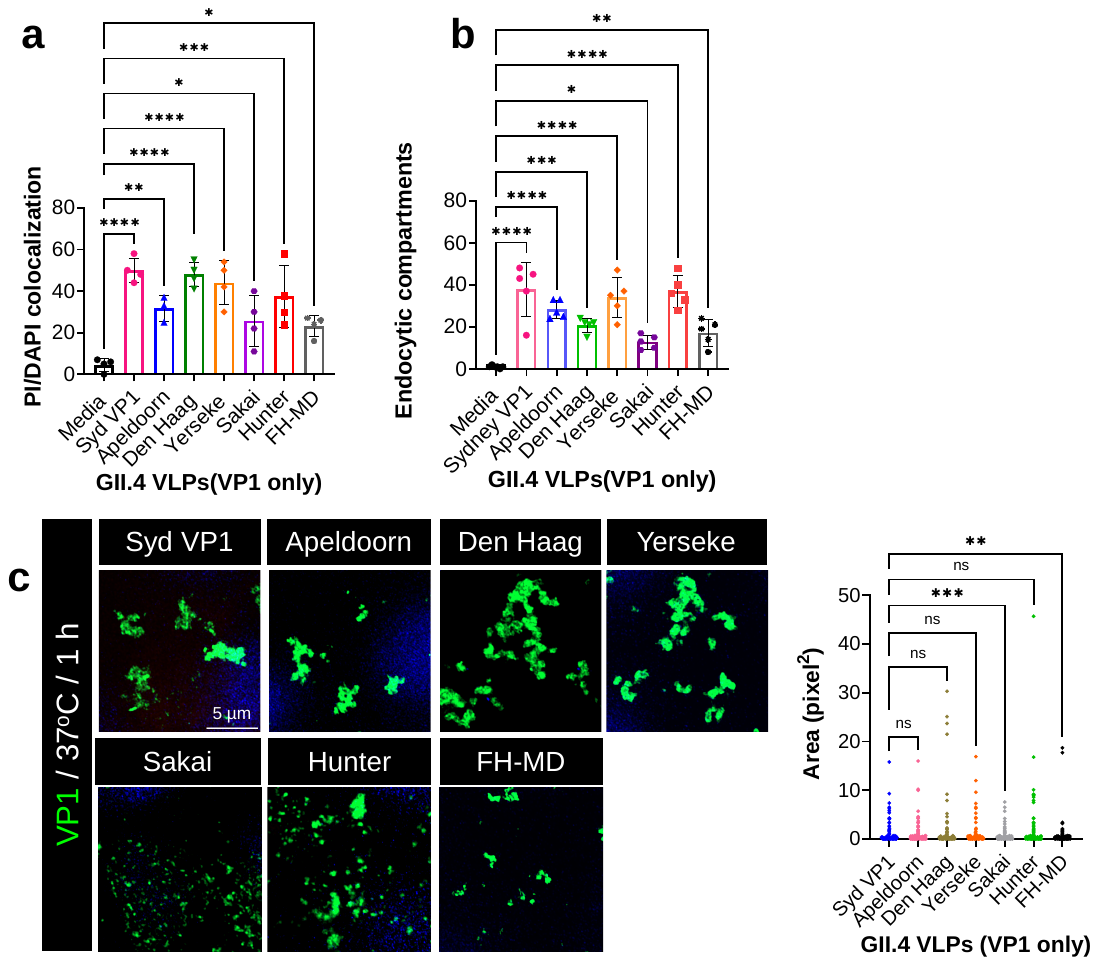

**Supplementary Figure 5. GII.4 HuNoV strains demonstrate variability in lysosomal exocytosis, endocytosis, and capsid clustering**

**(a)** VLP-induced membrane wounding calculated by quantifying DAPI/PI colocalization (ROI=4/condition). P values were calculated using one-way ANOVA, Dunnett’s Multiple Comparisons Test, where **** P ≤ 0.0001, *** P ≤ 0.001, ** P ≤ 0.01, * P ≤ 0.05 (n=3). **(b)** Quantitation of endocytosis calculated by FM-1-43X uptake induced by different GII.4 strains post 10 min VLP treatment. (ROI=5). P values (**** P ≤ 0.0001, *** P ≤ 0.001, ** P ≤ 0.01, * P ≤ 0.05) were calculated using one-way ANOVA, Dunnett’s multiple comparisons test (n=3). **(c)** Capsid clustering of GII.4 capsid (green) observed by incubating J2 HIEs with VLPs for 1 h at 37^o^C (n=2). Graph on the right represents the quantification of capsid clustering showing the difference between clustering (majority of clusters >5 pixel^2^, 1 pixel = 0.04 µm) vs non clustering variants (majority <1 pixel^2^).

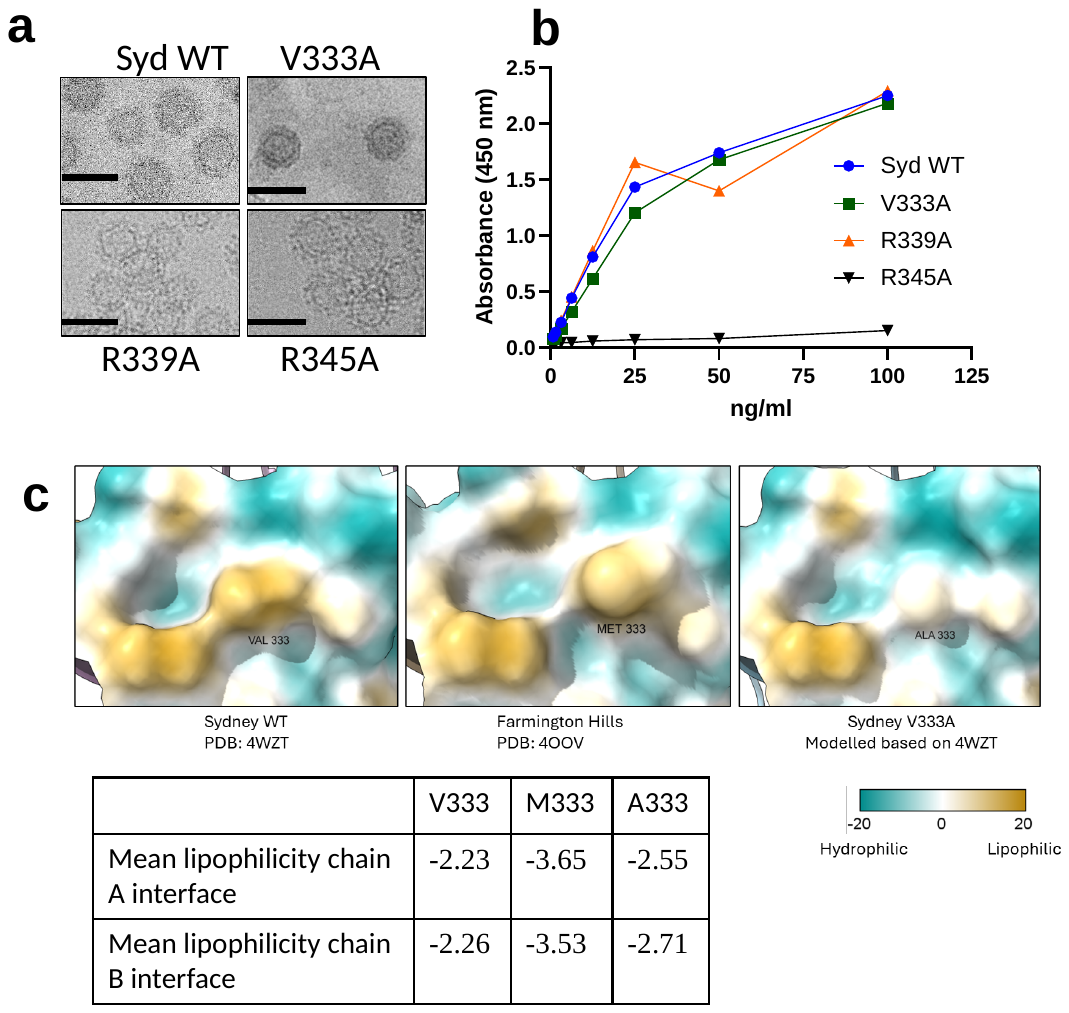

**Supplementary Figure 6. The capsids of GII.4 Syd mutants exhibit similar HBGA binding but exhibit diversity in their P dimer interface**

**(a)** Representative cryo-EM images of GII.4 Sydney and mutant VLPs collected on a Titan Krios instrument at 64,000 magnification. Black bars= 50 nm. **(b)** HBGA binding of GII.4 Sydney WT and mutants to PGM by ELISA. **(c)** Images from *in silico* hydrophilicity mapping highlighting the change in hydrophobicity around the P dimer interface between V333, M333 and A333 using ChimeraX v1.9. The table lists the lipophilicity of the interaction surfaces in each monomer of the P dimer, with lower values indicating an increase in hydrophilicity.
